# Fuzzy-FishNet: A highly precise distribution-free network approach for feature selection in clinical proteomics

**DOI:** 10.1101/024430

**Authors:** Wilson Wen Bin Goh

## Abstract

Network-based analysis methods can help resolve coverage and inconsistency issues in proteomics data. Previously, it was demonstrated that a suite of rank-based network approaches (RBNAs) provides unparalleled consistency and reliable feature selection. However, reliance on the t-statistic/t-distribution and hypersensitivity (coupled to a relatively flat p-value distribution) makes feature prioritization for validation difficult. To address these concerns, a refinement based on the fuzzified Fisher exact test, Fuzzy-FishNet was developed. Fuzzy-FishNet is highly precise (providing probability values that allows exact ranking of features). Furthermore, feature ranks are stable, even in small sample size scenario. Comparison of features selected by genomics and proteomics data respectively revealed that in spite of relative feature stability, cross-platform overlaps are extremely limited, suggesting that networks may not be the answer towards bridging the proteomics-genomics divide.

## Introduction

Mass spectrometry (MS)-based proteomics is now a key aspect of contemporary biological and clinical research. Although MS-based proteomics has advanced significantly in recent years, data reliability issues still persist. The standard setup is the Data-Dependent Acquisition (DDA) platform where eluting peptides from a separation column are selected for fragmentation semi-stochastically, leading to inconsistent quantitation amongst identified proteins, thus presenting a severe analytical challenge. Recent technological advancements have led to the new Data-Independent Acquisition (DIA) paradigm, where fragment precursor selection is independent of stoichiometry, leading to more spectral coverage (1, 2). An instance of DIA is SWATH, where data is captured by repeatedly cycling through precursor isolation windows (SWATH windows) of defined m/z ranges (3).

However, even with advanced techniques like SWATH, proteome coverage and signal quality issues persist. While Guo et al. have shown that, when coupled with PCT (Pressure Cycling Technology), SWATH could be used to reproducibly digitize the proteome of minute amounts of clinical samples in a high-throughput fashion (4), what was noteworthy, and of concern, was also the fact that SWATH is ostensibly noisier due to the concurrent fractionation of a large number of precursors.

Networks can be combined with proteomics synergistically to overcome its idiosyncratic data issues (5-10). For example, coverage and reliability of predictions can be improved dramatically simply using subnets (short for subnetworks) as contextualization (11-14). In particular, using the recently published clear cell renal cancer SWATH dataset of Guo *et al* (4), the efficacy of a suite of novel network-based analysis techniques termed Rank-Based Network Approaches (RBNAs) was demonstrated. Broadly, RBNAs work in the following steps: 1/ Features are ranked in inverse order (highest to lowest abundance) for each tissue. A cut-off at a predefined alpha level is used to identify the set of top alpha features for each tissue. 2/ Relevant features are used to fragment known pathways into subnets. Relevant features are the set of top-ranked protein defined subnets supported by a reasonable proportion amongst the samples within a class. Alternatively, where coverage is limited, a vector of known biological complexes can be used in its place. 3/ Each subnet is scored on each tissue according to the expression levels of the features constituting the subnet. Class-specific weights are introduced to modulate the scores. And finally, differential subnets are determined in the statistical feature-selection step. For details, refer to **Material and methods**.

Previously, it was shown that RBNAs have very high feature-selection stability and precision-recall rates, and work well in the small sample size scenario. The existing suite of RBNAs include SubNetworks (SNET), Fuzzy SNet (FSNET), Paired FSNet (PFSNET) and class-Paired PFSNet (PPFSNET). PFSNET and PPFSNET were the two best techniques, and performed very well on all performance benchmarks (14). However, there are two limitations worth investigating further --- 1/ feature selection based on the modified t-statistic and t-distribution may not be a valid assumption and 2/ PFSNET and PPFSNET tend to be extremely sensitive, making a fairly large number of predictions for which the p-value distribution is relatively flat (many of these are 0 or close to 0), this makes it difficult to prioritize which subnets to test and validate first.

To deal with these problems, a new addition to the RBNA family is introduced --- Fuzzy-FishNet. Fuzzy-FishNet uses a weighted version of the Fisher’s exact test to derive an exact probability for whether a subnetwork is differentially expressed between the normal and cancer classes. We demonstrate the efficacy (based on precision-rate and feature selection stability) of Fuzzy-FishNet to its non-weighted counterpart FishNet, the standard single protein-based two sample t-test (SP), hypergeometric enrichment (hypgeo or HE), and against the existing RBNAs. We also compared the networks predicted by proteomic and genomic data for clear cell renal cancer to investigate if network-based analysis can give rise to improved correlations.

## Material and methods

### SWATH data

The SWATH dataset of Guo *et al* was used in this study (4). This dataset contains 24 SWATH runs from 6 pairs of non-tumorous and tumorous clear-cell renal carcinoma (ccRCC) tissues, which have been swathed in duplicates (12 normal, 12 cancer).

### SWATH data interpretation

All SWATH maps were analyzed using OpenSWATH (15) and a spectral library containing 49,959 reference spectra for 41,542 proteotypic peptides from 4,624 reviewed SwissProt proteins (4). The library was compiled using DDA data of the kidney tissues in the same mass spectrometer. Protein isoforms and protein groups were excluded from this analysis. The peptides identified were aligned prior to protein inference using the algorithm TRansition of Identification Confidence (TRIC) (version r238), which is available from https://pypi.python.org/pypi/msproteomicstools and https://code.google.com/p/msproteomicstools. The parameters used for the feature_alignment.py program are: max_rt_diff=30, method=global_best_overall, nr_high_conf_exp=2, target_fdr=0.001, use_score_filter=1. The two most intense peptides were used to quantify proteins. 3,123 proteins were quantified across all samples with peptide and protein FDR below 1%.

### Next-generation sequence data (genomics)

For comparison, a genomics (Illumina HiSeq 2000 RNA sequencing platform) dataset for clear cell renal cancer is derived from The Cancer Genome Atlas (https://tcga-data.nci.nih.gov/docs/publications/kirc_2013/).

The dataset comprises 31 normal samples and 31 tumor samples covering 18,400 genes (16). Gene expression was quantified by counting the number of reads overlapping each gene model’s exons and converted to Reads per Kilobase Mapped (RPKM) values via division by the transcribed gene length.

### Protein complexes (Subnets)

Subnets can be determined a priori and independent of the data used for analysis. For example, decomposition of a network into subnets can be optimized via functional coherence evaluation (11). However, protein complexes are true biological subnets, and shown to be superior to inferred ones (13). They are also stable as they are determined independently of the experimental data. Thus the complex-based feature vector can be used in generalizability studies comparing related genomic and proteomic data.

Protein complexes were obtained from CORUM database which contains manually annotated protein complexes from mammalian organisms (17). Complexes with at least 3 proteins that were identified and measured in the proteomics screen were retained (1363 complexes).

### Standard protein-based feature selection using t-test (SP)

As a control to why network methods are required to extend proteomic analysis, a t-statistic (*T*_*p*_) is calculated for each protein p by comparing the z-normalized expression scores between classes *C*1 and *C*2, with the assumption of unequal variance between the two classes (18).

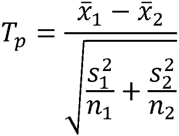

where 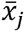 is the mean expression level of the protein *p*, *s*_*j*_ is the standard deviation and *n*_*j*_ is the sample size, in class *Cj*.

The *T*_*p*_ is compared against the nominal t-distribution to calculate the corresponding p-value. A feature is deemed significant if p-value ≤ 0.05.

### Hypergeometric Enrichment (HE)

HE is a standard hypergeometric enrichment pipeline performed in many earlier studies (6) and consists of two parts: 1/ Differential proteins are identified using the unpaired two-sided t-test between normal and disease samples using their z-normalized protein expressions (This is similar to SP) (19). Proteins with p-value ≤ 0.05 are considered differential. 2/ This is followed by a hypergeometric enrichment analysis against the protein complexes (p-value ≤ 0.05). Given a total number of proteins *N*, with *B* of these belonging to a complex and *n* of these proteins in the test set, the probability *P* that *b* or more proteins from the test set are associated by chance with the complex is given by:

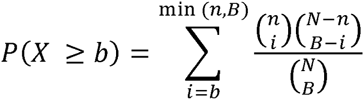

The complex is deemed significant in HE if P(*X* ≥ *b*)≤ 0.05.

### Fuzzy-FishNet/FishNet

Fuzzy-FishNet and FishNet are similar methods differing only in weights assigned to the rank proteins. Fuzzy-FishNet uses rank weights while FishNet uses a binary metric (see below).

We begin with a description of FishNet: Given a protein *gi* and a tissue *pk*, let fs(*gi,pk*) = 1, if the protein *gi* is among the top alpha percent (default = 10%) most-abundant proteins in the tissue *pk*; and = 0 otherwise.

For a complex S, and samples in class J, C_j_, and samples in class k, C_k_. We can express the distribution of proteins in the top alpha percent between C_j_ and C_k_ against S in a contingency table as:

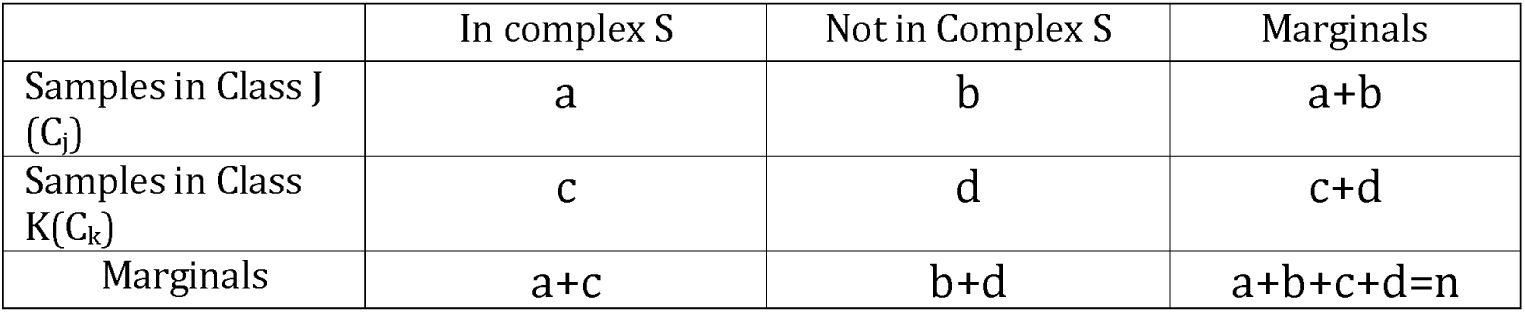

|  | In complex S | Not in Complex S | Marginals |
| --- | --- | --- | --- |
| Samples in Class J ($C_j$) | a | b | a+b |
| Samples in Class K ($C_k$) | c | d | c+d |
| Marginals | a+c | b+d | a+b+c+d=n |

where a and c are the sum of alpha proteins for samples in class C_j_ and C_k_ mappable to proteins within complex S respectively, and b and d are the sum of proteins across samples in class C_j_and C_k_ that are missed for proteins in complex S respectively.

The fisher exact probability p of obtaining this given set of values is then:

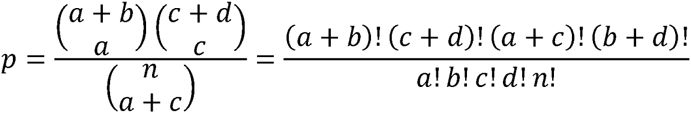

The fisher exact probability is also the hypergeometric probability of observing this particular arrangement of the data, assuming the given marginal totals, on the null hypothesis that both C_j_ and C_k_ have similar distributions of alpha proteins across their class members mappable to proteins in complex S.

To calculate a significance value (p-value) for a given observed probability p, we can sum the probabilities of obtaining a value equal to or greater than the observed p in a one-sided test. Alternatively, when *n* is relatively large, and a, b, c and d are all greater than 5, the p-value for the observed fisher exact probability can be approximated using the chi-square distribution. Given that the fisher test is known to be conservative — i.e., its actual rejection rate is lower than the nominal level, and that the actual data distribution might not match well theoretical distributions, we rank p in order of increasing value, and select the top 1% features.

For Fuzzy-FishNet, the definition of the function fs(gi,pk) is replaced so that fs(gi,pk) is assigned a value between 5 and 0 as follows: fs(gi,pk) is assigned the value 5 if gi is among the top alpha1 percent (default = 10%) of the most-abundant proteins in pk. It is assigned the value 0 if gi is not among the top alpha2 percent (default = 20%) most-abundant proteins in pk. The range between alpha1 percent and alpha2 percent is chopped into *n* equal-sized bins (default =4), and fs(gi,pk) is assigned the value 4, 3, 2, or 1 depending on which bin gi falls into in pk. As with FishNet, the top 1% of features is selected.

### SNET/FSNET/PFSNET/PPFSNET

The RBNAs (SNET, FSNET, PFSNET and PPFSNET) are similar algorithms but differing in certain key assumptions or test set-ups.

We begin with a description of SNET:

Given a protein *gi* and a tissue *pk,* let *fs(gi,pk)* = 1, if the protein *gi* is among the top alpha percent (default = 10%) most-abundant proteins in the tissue *pk*; and = 0 otherwise.

Given a protein *gi* and a class of tissues *Cj*, let

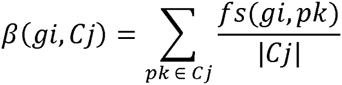

That is, *β*(*gi, Cj*) is the proportion of tissues in *Cj* that have *gi* among their top alpha percent most-abundant proteins.

Let score(*S,pk,Cj*) be the score of a protein complex *S* and a tissue *pk* weighted based on the class *Cj*. It is defined as:

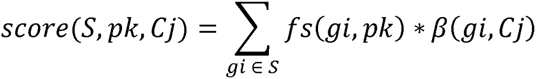

The function *f*_*SNET*_ (*S, X, Y, Cj*) for some complex *S* is a t-statistic defined as:

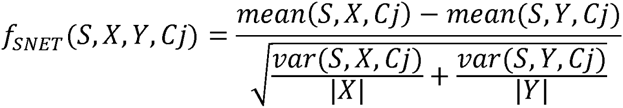

where *mean*(*S*,#,*Cj*) and *var*(*S*,#,*Cj*) are respectively the mean and variance of the list of scores {*score*(*S,pk,Cj*) | *pk* is a tissue in # }.

The complex *S* is considered significantly highly abundant (weighted based on *Cj*) in *X* but not in *Y* if *f*_*SNET*_(*S,X,Y,Cj*) is at the largest 5% extreme of the Student t-distribution, with degrees of freedom as determined by the Welch-Satterwaite equation.

Given two classes *C*1 and *C*2, the set of significant complexes returned by SNET is the union of {*S* | *f*_*SNET*_(*S,C1,C2,C1*) is significant} and {*S* | *f*_*SNET*_(*S,C2,C1,C2*) is significant}, the former being complexes that are significantly consistently highly abundant in *C1* but not *C2*, the latter being complexes that are significantly consistently highly abundant in *C2* but not *C1*.

FSNET is identical to SNET, except in one regard:

For FSNET, the definition of the function *fs*(*gi,pk*) is replaced so that *fs*(*gi,pk*) is assigned a value between 1 and 0 as follows: *fs*(*gi*,*pk*) is assigned the value 1 if *gi* is among the top alpha1 percent (default = 10%) of the most-abundant proteins in *pk*. It is assigned the value 0 if gi is not among the top alpha2 percent (default = 20%) most-abundant proteins in pk. The range between alpha1 percent and alpha2 percent is chopped into *n* equal-sized bins (default =4), and *fs*(*gi,pk*) is assigned the value 0.8, 0.6, 0.4, or 0.2 depending on which bin *gi* falls into in *pk*.

A test statistic *f*_*FSNET*_ is then defined analogously to *f*_*SNET*_. Given two classes *C1* and *C2,* the set of significant complexes returned by FSNET is the union of {*S* | *f*_*FSNET*_(*S,C1,C2,C1*) is significant} and {*S* | *f*_*FSNET*_(*S,C2,C1,C2*) is significant}.

For PFSNet, the same *fs*(*gi,pk*) function as in FSNet is used. But it defines a score delta(*S,pk,X,Y*) for a complex *S* and tissue *pk* wrt classes *X* and *Y* as the difference of the score of *S* and tissue *pk* weighted based on *X* from the score of *S* and tissue *pk* weighted based on *Y*. More precisely: *delta*(*S,pk,X,Y*) = *score*(*S,pk,X*)– *score*(*S,pk,Y*).

If a complex *S* is irrelevant to the difference between classes *X* and *Y*, the value of *delta*(*S,pk,X,Y*) is expected to be around 0. So PFSNet defines the following one-sample t-statistic:

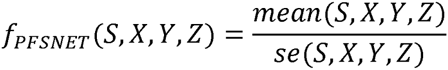

where mean(*S, X, Y, Z*) and *se*(*S, X, Y, Z*) are respectively the mean and standard error of the list {*delta*(*S, pk, X, Y*) | *pk* is a tissue in *Z*}. The complex *S* is considered significantly consistently highly abundant in *X* but not in *Y* if *f*_*PFSNet*_(*S, X, Y, X∪ Y*) is at the largest 5% extreme of the Student t-distribution.

Given two classes *C1* and *C2*, the set of significant complexes returned by PFSNet is the union of {*S* | *f*_*PFSNet*_(*S,C1,C2,C1*∪ *C2*) is significant} and {*S* | *f*_*PFSNet*_(*S,C2,C1,C1*∪ *C2*) is significant}, the former being complexes that are significantly consistently highly abundant in *C1* but not *C2,* the latter being complexes that are significantly consistently highly abundant in *C2* but not *C1*.

The above formulation of PFSNet is for the situation where tissues in *C1* and *C2* are unpaired. If paired tissues are used, a paired-sample version of PFSNet (PPFSNET) can be formulated as follows.

Given a subject *pk*, we write *pkA* to denote his tissue in class *C1* and *pkB* to denote his paired tissue in class *C2*. Then we define the following paired delta score of the complex *S* and subject *pk* wrt classes *X* and *Y*:

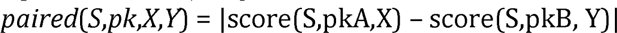

If the complex *S* is irrelevant to the difference between classes *X* and *Y*, as mentioned earlier, then the mean of *paired*(*S,pk,X,Y*) is expected to be 0. We define a one-sample t-statistic to test for this:

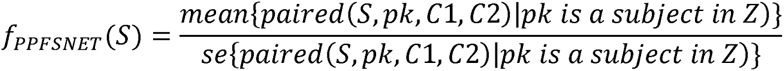

where *Z* is the set of all subjects with paired tissues in *C1* and *C2*. A complex S is considered significant, if f_PPFSNET_(S) is at the largest 1% extreme of the Student t-distribution.

### Performance Benchmarks

Precision/recall and feature-selection stability were used as performance benchmarks. Cross-validation predictive accuracy is omitted since FishNet does not provide feature-level scores per sample.

#### Precision/Recall

In precision/recall, the significant complexes *c*, from each subsampling simulation is benchmarked against the total set of significant complexes, *C*, derived from an analysis of the complete dataset. We make the assumption that the complete dataset is representative of the population. Thus, a completely precise method based on a subsampling should report a subset *c* of *C* (*c* ⊆ *C*) as significant, and no more (considered false positives). Similarly, perfect recall should report all complexes in *C* (i.e., *c* = *C*) as significant.

Precision and recall are calculated as follows:

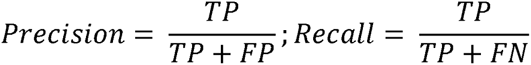

where TP, FP and FN are the True Positives, False Positives and False Negatives respectively.

Both measurements are important in evaluating the performance of a method. A method that is precise but not sensitive would make some good-quality predictions but may not provide enough data for model building or understanding the phenomena whereas a highly imprecise but sensitive method may capture all relevant features but at the cost of introducing much noise (irrelevant features). A good method must be both precise and sensitive (high recall). To evaluate both precision and recall concurrently, we can use the F-score (*Fs*), which is basically the harmonic mean:

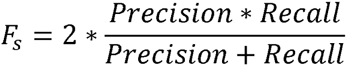

These evaluation metrics require prior knowledge of the set of TP, TN, FP and FN in the dataset. In biology and particularly in proteomics, such “gold standard” data does not exist. We therefore make the assumption that the full dataset is the population, and apply the given feature-selection algorithm to determine the total set of TPs. Comparisons of features selected by repeated subpopulation sampling against those from the complete dataset provides an estimate of the precision and recall rates. However, we must highlight a caveat to this way of defining the gold standard: It may mislead when the given feature-selection algorithm is unstable, as the algorithm is likely to return an entirely different “gold standard” set of features when applied to a different dataset.

**Feature-selection stability** — A good feature-selection method must be able to make consistent and reproducible selections, even at small sample sizes. Across different samplings, the technique should reliably provide similar findings. A method with generally high accuracy but low stability has limited utility. It is well known that depending on the dataset, or different parts of the dataset, the same test can select highly different feature sets; this can be attributed to a lack of statistical power and/or unreliable p-value.

For each method, we took random samplings of size 4, 6 and 8 tissues from both normal and cancer classes (n=12) to simulate small (4) to moderate (8) sample-size scenarios. This is repeated 1,000 times to generate a binary matrix, where each row is a simulation, a value of 1 indicates a complex is significant, and 0 otherwise.

The binary matrix is used for comparing stability and consistency of significant features produced by each method. Two evaluations on the binary matrix were performed: 1/ row-wise comparisons based on the Jaccard coefficient to evaluate feature vector pairwise similarity, and 2/ column summation to evaluate the persistence/stability of each selected significant complex.

The feature-selection stability score is calculated as follows: The columns in the binary matrix generated per bootstrap sampling represent the all complexes being tested, while the rows represent the number of simulations (n=1,000). A value of 1 means the complex turned up significant while 0 means it did not. Summing each column and dividing it by the number of simulations provides a single stability vector containing the normalized values indicative of complex stability (0 means the complex was never observed, while 1 means the complex was significant across all 1,000 simulations). To calculate a unified score for feature-selection stability, first, all 0 values are discarded from the stability vector (since these are complexes that have never been observed even once across all simulations, and thus irrelevant). Next, the remaining values are summed and divided by the total length of the stability vector, thus generating the feature-selection stability score.

## Results and discussions

### Fuzzy-FishNet addresses over-reliance on the t-distribution and limits feature-selection inflation

The design for Fuzzy-FishNet stems from the need to address two potential flaws in earlier RBNAs: the first being the assumption that the data has to approximate a t-distribution (All current RBNAs use the t-statistic, and compares it to a reference t-distribution), and the second being the need to reduce the number of features being predicted (limiting over-sensitivity), while maintaining excellent precision-recall.

The Fisher’s test has some useful properties for the first problem. First, it is valid for small sample sizes, which is fairly common when dealing with biological data (when *n* ≤ 5). Secondly, it is an exact test, which means that the extent of deviation from a null hypothesis can be calculated determinably, rather than reliance on a theoretical distribution (as well with the t-test).

In the second concern, as mentioned, while the most powerful RBNAs (PFSNET and PPFSNET) exhibit very high feature-selection stability, as well as precision-recall rates (14), it should be noted they also report a relatively large number of features as well. Significant features selected by PFSNET and PPFSNET have a rather flat p-value distribution i.e., many of the features had p-values at 0 or close to 0 (Supplementary Data 1) thus making it difficult to prioritize which features to test and validate experimentally. In FishNet and Fuzzy-FishNet, because exact probabilities can be calculated, it should generate values with relative high resolution, allowing the features to be ranked. Furthermore, to limit the number of features selected, instead of returning all features that meet some statistical threshold cut-off (e.g. 0.05 or 0.01), we ranked the fisher exact probabilities calculated for each complex from lowest to highest, and selected the top 1% (1363*0.01≈14). The downside to this procedure is that regardless, 14 features will always be selected even if these are not relevant. However, the stability of the ranks of these features can be checked given the assertion that if these features are meaningful, then they should be repeatedly observed (at the top 1%) given tests on any random subset of the full data. Otherwise, the ranks would be highly unstable.

### The fuzzification procedure in Fuzzy-FishNet selects better quality features

The fuzzification procedure in Fuzzy-FishNet is inherited from methods such as FSNET, PFSNET(20) and PPFSNET(14). The purpose of fuzzification is to weigh the signal more favorably from proteins that are highly ranked, while allowing signal from lower ranked proteins to also be included, thus boosting sensitivity.

In FSNET/PFSNET/PPFSNET, the weights are interpolated from 1 to 0 as continuous variables. In Fuzzy-FishNet, the weights have to be integers due to the permutation-based calculations. To determine if fuzzification improves the quality of feature selection, we compared Fuzzy-FishNet to a non-fuzzified variant, FishNet.

Figures 1A and 1B shows the CORUM Complex IDs, contingency tables and Fisher probability for the top 5 complexes in Fuzzy-FishNet and FishNet respectively. The top 3 complexes are similar, but it can also be seen that due to fuzzification, the p for Fuzzy-FishNet is smaller — i.e., harder to observe the data distribution by chance. It is noteworthy that that most complexes have high p (Supplementary Figure 1). There is deep overlap of complexes between Fuzzy-FishNet and FishNet (Figure 1C). Interestingly, the complement showed that the 5 Fuzzy-FishNet only complexes corresponded to only 17 proteins while the 5 FishNet only complexes corresponded to a large 109 proteins. This suggests that the 14 complexes in Fuzzy-FishNet are more homogeneous than in FishNet. The 5 were probably missed because these complexes are smaller and/or the signal is weaker (e.g. there are fewer overlapping proteins but these are actually highly ranked). Obviously, the signal can be accentuated by the fuzzification procedure.

**Figure 1.**
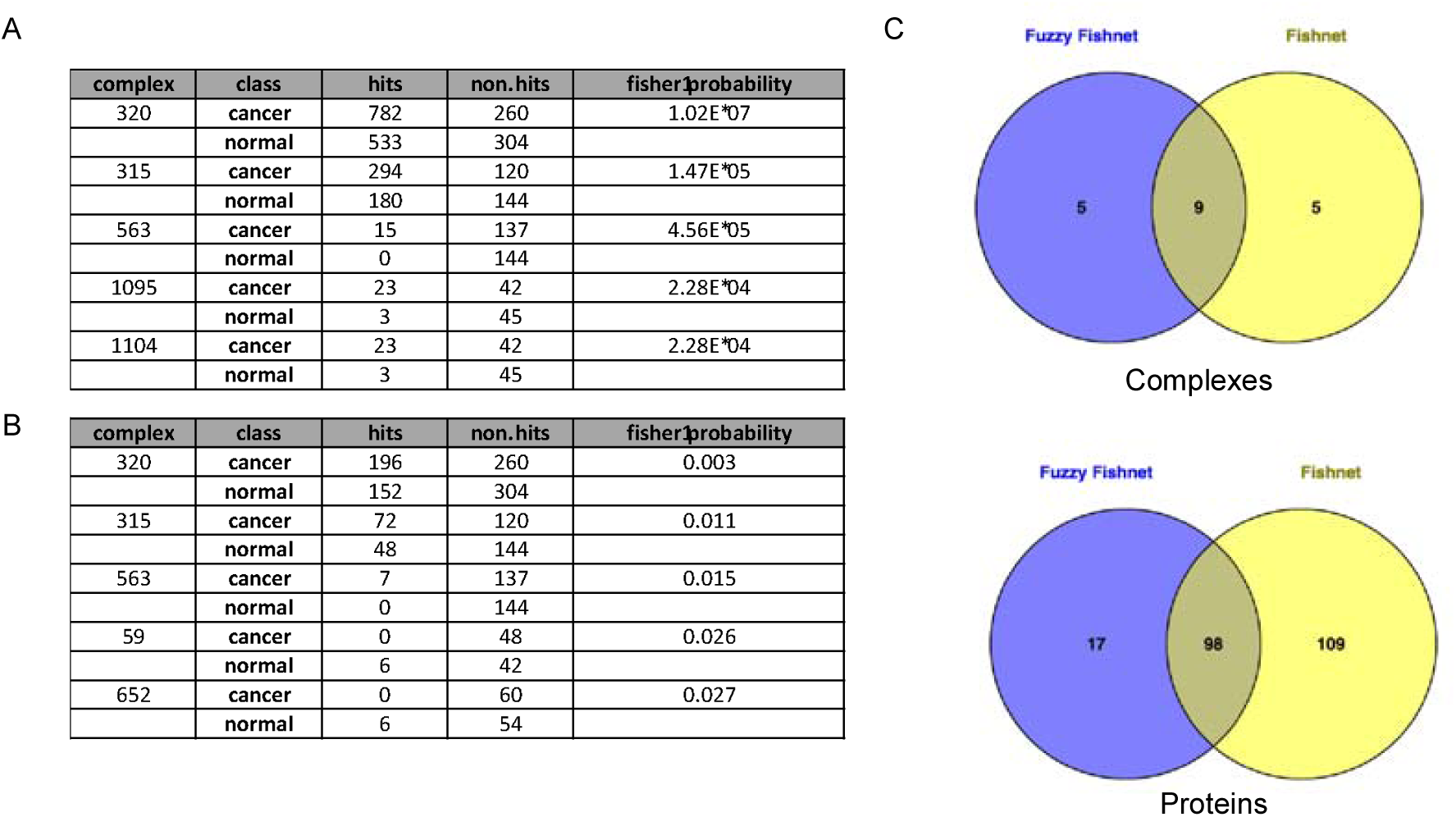
Comparison of significant complexes and overlaps between Fuzzy-FishNet and FishNet. **A: Top 5 complexes selected by Fuzzy-FishNet.** The table shows the CORUM IDs for the complex, the contingency table and the corresponding Fisher exact probability that was calculated. **B: Top 5 complexes selected by FishNet**. As before, the table shows the CORUM IDs for the complex, the contingency table and the corresponding Fisher exact probability that was calculated. Note that the top three complexes are similar (c.f. Figure 1A). **C: Complex and protein overlap between Fuzzy-FishNet and FishNet.** There is deep sharing of complexes between Fuzzy-FishNet and FishNet. Interestingly, the complement show that the 5 Fuzzy-FishNet only complexes corresponded to only 17 proteins while the 5 FishNet only complexes corresponded to a large 109 proteins. This result shows that the 14 complexes in Fuzzy-FishNet are more homogeneous. The 5 were probably missed because they are smaller and/or the signal is weaker. Obviously, the signal can be accentuated by the fuzzification procedure.

To confirm if the selected complexes could discriminate sample classes, we derived the constituent protein expressions from selected complexes (115 for Fuzzy-FishNet and 207 for FishNet), and performed hierarchical clustering (Euclidean Distance, Ward’s linkage). Figure 2 shows that for the large part, normal and cancer classes can be discriminated using either methods. However the class segregation for Fuzzy-FishNet is stronger. Normal samples 6, 7 and 8 from the second replicate are consistently misgrouped with the cancer branch. Using other analysis methods, this misgrouping was also observed (21).

**Figure 2.**
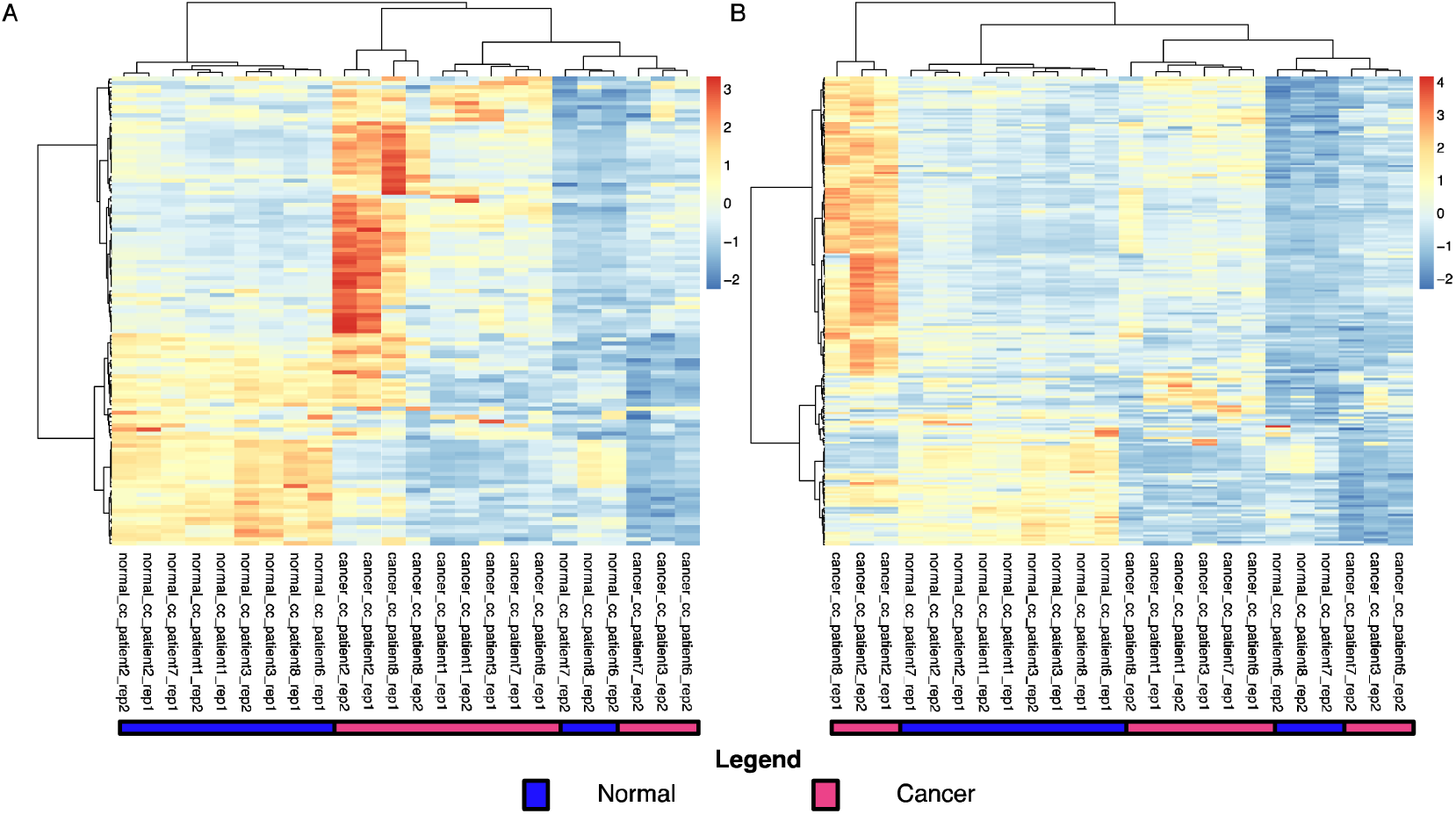
Hierarchical clustering (HCL) of proteins (from significant complexes) for Fuzzy-FishNet and FishNet. A: HCL (Fuzzy-FishNet). The tree (Euclidean Distance, Ward’s linkage) shows relatively better separation than the corresponding tree for FishNet (c.f. Fig 2B). **B: HCL (FishNet).** The tree (Euclidean Distance, Ward’s linkage) shows relatively better separation than the corresponding tree for FishNet (c.f. Fig 2B). **B: HCL (FishNet).** The tree (Euclidean Distance, Ward’s linkage) shows relatively poorer separation than the corresponding tree for FishNet (c.f. Fig 2A). This provides some indication that the complexes selected by Fuzzy-FishNet are more informative. Note that normal samples 6,7,8 from replicate 2 are consistently misclassified.

Feature-selection stability, pairwise feature vector similarity and false positive rates are compared between Fuzzy-FishNet, FishNet, the standard single protein t-test (SP), and the hypergeometric test (hypgeo) (Figure 3). Random selection of 4, 6 and 8 samples from each class was performed 1000 times to evaluate how persistent each feature is (Figure 3A), and how similar pairwise samplings were (Figure 3B). Amongst these, Fuzzy-FishNet demonstrated that it was able to make very reliable predictions, and appeared to be fairly robust even at small sample sizes. Moreover, it was able to achieve these with the lowest false positves rates as well (Figure 3C). This suggests that the rank order amongst the top 1% is conserved.

**Figure 3.**
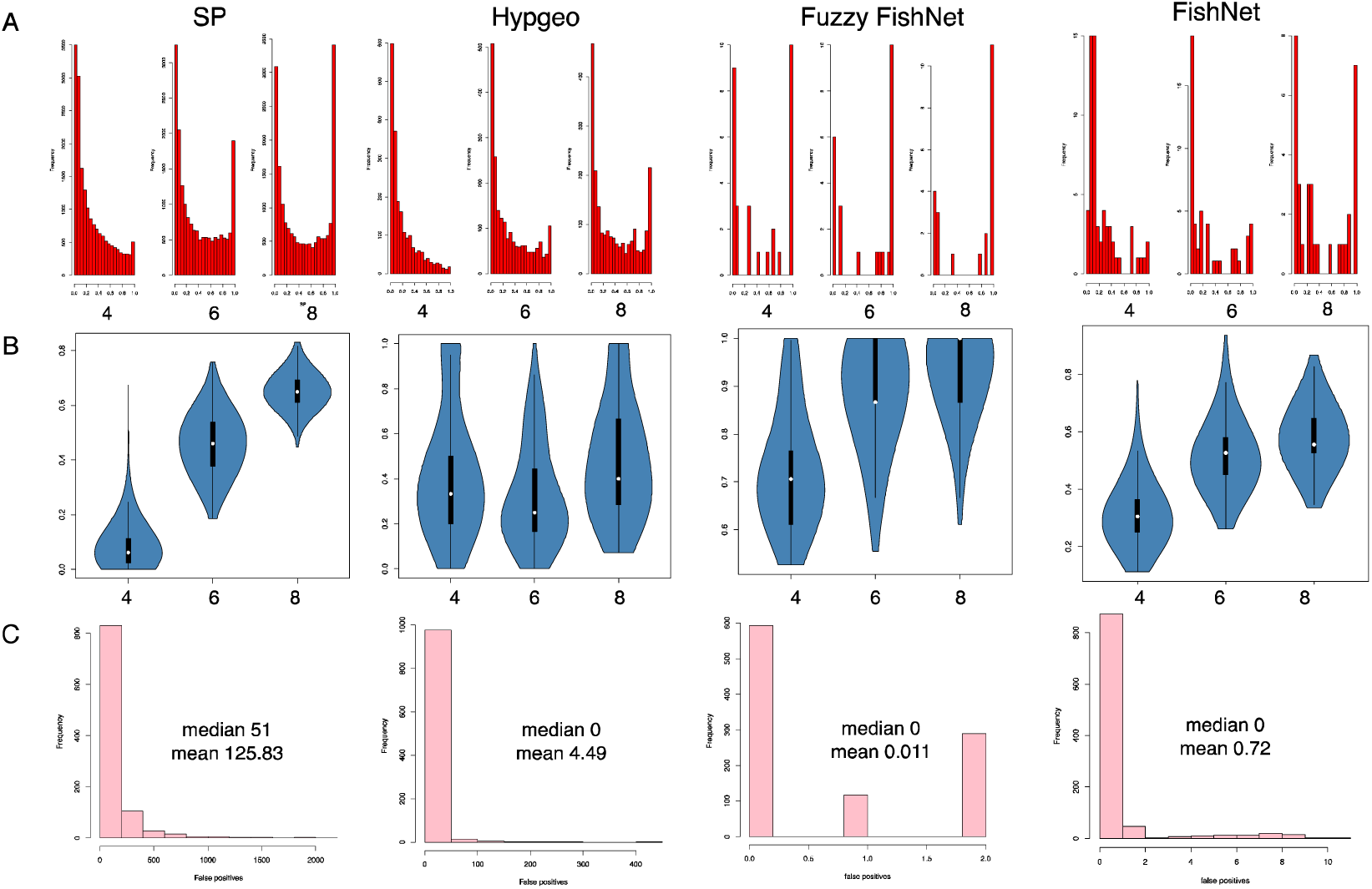
Performance metrics (Feature-selection stability, pairwise feature vector similarity and false positive distribution) for single protein standard t-test (SP), Hypergeometric Enrichment (Hypgeo), Fuzzy-FishNet and FishNet. A: Feature-selection stability across 1,000 simulations. With increased sampling size (from 4 to 8), SP, HE and FishNet’s feature-selection stability improved, i.e., an observable right shift in the histograms. In these cases however, a vast majority of selected features was never consistently reproduced. Fuzzy-FishNet responded well to sample size increments, with an obvious right-shift indicating many of the features were stably observed. **B: Pairwise feature vector similarity across random samplings.** SP, HE, Fuzzy-FishNet and FishNet were evaluated 1,000 times on random subsets of sizes 4, 6 and 8. Simulations were compared pairwise for reproducible features using the Jaccard Coefficient. Fuzzy-FishNet excelled here, even in small sample size scenario. **C: False-positive distribution across 1,000 simulations**. Samples from the normal class were randomly assigned to two groups, with feature selection performed using each method. Fuzzy-FishNet has the lowest false positive rate.

To further determine if the selected features are meaningful, we calculated the precision-recall rates from random subsets of the data and compared it against features selected in the full dataset (Table 1). Fuzzy-FishNet excelled in both precision and recall with the highest F-scores amongst the methods tested. It should be noted that the high precision observed for SP is inflated. To examine this, the p-value threshold was adjusted to restrict the number of allowed features in SP to approximately the top 500. Noticeably, by restricting the number of features, there is a drop in the proportion of stable features (Supplementary Figure 2A compared to Figure 1A; SP) which suggests substantial fluctuations in the ranks of those features which met the original threshold requirement (p-val ?0.01). This is accompanied by a concomitant drop in the pairwise feature similarity (Supplementary Figure 2B). Supplementary Figure 2C shows that the increased feature-selection stringency produces a drastic drop in recall while precision is maintained. These results show that the perceived stability and consistency produced by SP is likely an artifact due to the large number of features it reports.

**Table 1.** **Precision-Recall performance for Fuzzy-FishNet (A), FishNet (B), standard t-test (sp) and hypergeometric-enrichment (hypgeo) (C).** Fuzzy-FishNet performs extremely, excelling both in precision and recall. It is noteworthy that it also functions very well in the small-sample size scenario.

|  |  |  |  |  |  |  |
| --- | --- | --- | --- | --- | --- | --- |
| A |  | **precision** | **recall** | **F-score** | **sd_precision** | **sd_recall** |
|  | **f_fishnet_4** | 0.891 | 0.909 | 0.900 | 0.087 | 0.073 |
|  | **f_fishnet_6** | 0.935 | 0.945 | 0.940 | 0.072 | 0.057 |
|  | **f_fishnet_8** | 0.961 | 0.967 | 0.964 | 0.058 | 0.044 |
| B |  | **precision** | **recall** | **F-score** | **sd_precision** | **sd_recall** |
|  | **fishnet_4** | 0.630 | 0.667 | 0.648 | 0.159 | 0.106 |
|  | **fishnet_6** | 0.784 | 0.794 | 0.789 | 0.083 | 0.080 |
|  | **fishnet_8** | 0.837 | 0.850 | 0.844 | 0.074 | 0.067 |
| C |  | **precision** | **recall** | **F-score** | **sd_precision** | **sd_recall** |
|  | **sp_4** | 0.916 | 0.455 | 0.608 | 0.068 | 0.073 |
|  | **sp_6** | 0.933 | 0.639 | 0.759 | 0.051 | 0.066 |
|  | **sp_8** | 0.933 | 0.639 | 0.759 | 0.051 | 0.066 |
|  | **hypgeo_4** | 0.211 | 0.181 | 0.195 | 0.204 | 0.198 |
|  | **hypgeo_6** | 0.389 | 0.477 | 0.428 | 0.172 | 0.201 |
|  | **hypgeo_8** | 0.477 | 0.463 | 0.470 | 0.142 | 0.245 |

### Fuzzy-FishNet selected features are supported by other RBNAs

The features selected by Fuzzy-FishNet, FishNet, PFSNET and PPFSNET are compared using a four-way Venn diagram (Figure 4A). All 9 intersecting complexes between FishNet and Fuzzy-FishNet were also reported by PFSNET and PPFSNET. The RBNAs also reported the Fuzzy-FishNet complement and FishNet complement as significant. Note that PFSNET and PPFSNET reports an additional 54 complexes.

**Figure 4.**
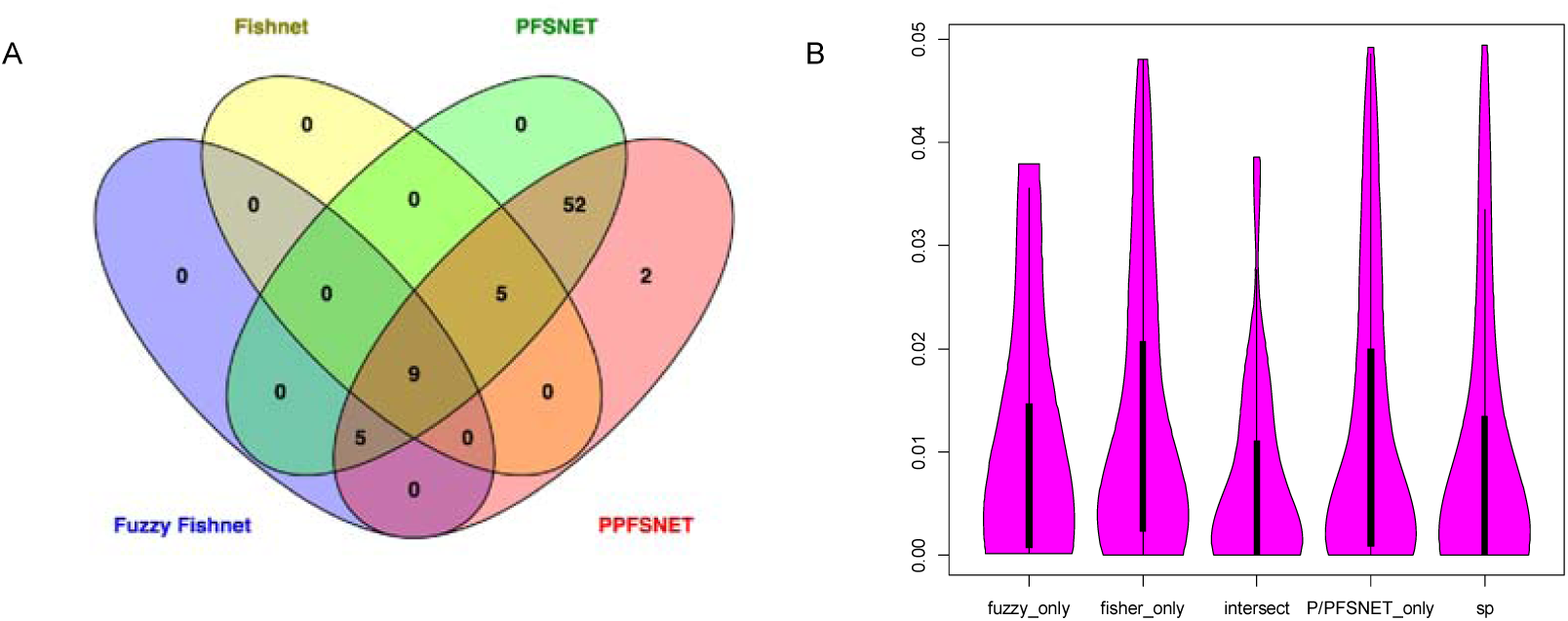
Comparisons of Fuzzy-FishNet against other network algorithms A: Overlaps between 4 rank-based network approaches (RBNAs): FishNet, Fuzzy-FishNet, PFSNET and PPFSNET. We compared the complexes predicted by FishNet and Fuzzy-FishNet against two recently developed RBNAs (PFSNET and PPFSNET). All complexes selected by FishNet and Fuzzy-FishNet were also selected by PFSNET/PPFSNET. However, PFSNET/PPFSNET picked up many additional complexes. Because the PFSNET/PPFSNET p-value distribution is generally flat, with most features with p-values at 0 or close to 0, these algorithms might be too sensitive, and prevent discrimination or prioritization of which complexes/proteins to test first. **B: Fuzzy-FishNet selects more significant t-test features**. Because the p-value distribution for PFSNET and PPFSNET lacks variability, we compared the standard t-test p-values of proteins (from 9 significant complexes) overlapped between FishNet and Fuzzy-FishNet (intersect), proteins found in FishNet only (fisher_only; from 5 complexes), proteins found in Fuzzy-FishNet only (fuzzy_only, 5 complexes), PFSNET and PPFSNET only complexes (54 complexes), and the remaining significant single protein t-test proteins (sp). The intersection and Fuzzy-FishNet only median points were lower, suggesting selection of highly significant proteins. On the other hand, the median of the FishNet only proteins were higher, suggesting selection of less significant proteins. Fuzzification improves the signal for relevant complexes that might otherwise be missed.

Since PFSNET and PPFSNET’s p-value distribution are both quite flat, we cannot say if the FishNet and Fuzzy-FishNet selected features are enriched for higher quality selections. As a getaround, this can be tested indirectly by comparing the distribution of t-test p-values for proteins found in significant complexes. Figure 4B shows that the intersect for Fuzzy-FishNet and FishNet, and Fuzzy-FishNet only are enriched for more significant proteins than FishNet only and PPFSNET/PFSNET only. Hence, although it reports fewer complexes than current generation RBNAs, these are likely to be higher quality.

Table 2 shows the feature-stability scores and F-scores (precision-recall) comparing the FishNet methods, all RBNAs, SP (single-protein t-test) and HE (Hypergeometric Enrichment). FishNet has very strong precision-recall, comparable to PPFSNET’s. Feature-selection stability however, is relatively weaker compared to P/PFSNET’s. However, it should be noted that P/PFSNET are likely hyper-sensitive, hence, it will tend to report similarly large sets of complexes regardless of sampling. Unfortunately, because the p-value distribution for P/PFSNET is quite flat, we cannot test the rank stability of the top 14 significant complexes, and compare these statistics directly against the Fisher-based methods.

**Table 2.**
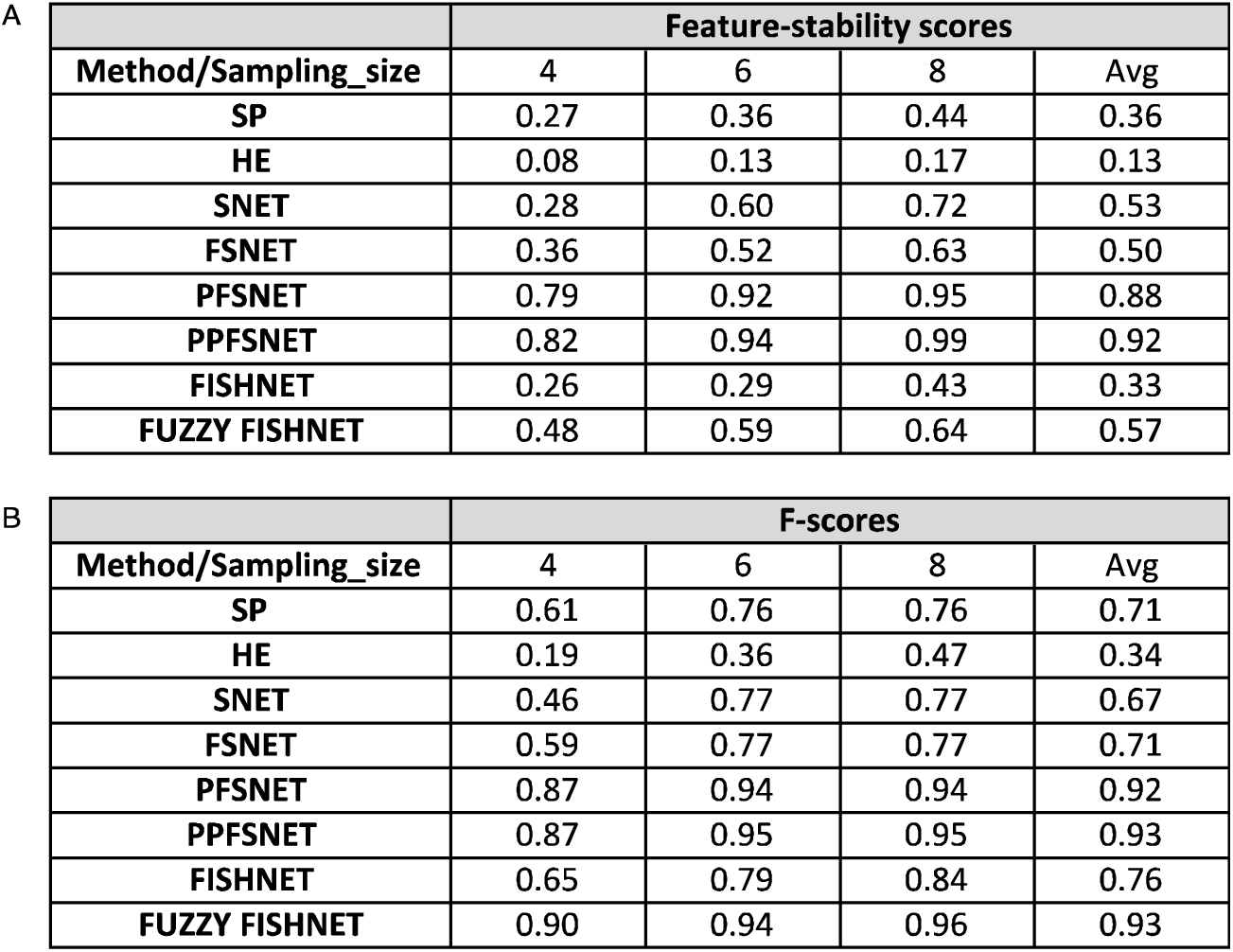
**Comparison of feature-stability scores (A) and F-scores (B) between standard t-test (SP), hypergeometric-enrichment (HE), the rank-based network analysis methods, SNET, FSNET, PFSNET and PPFSNET, FishNet and Fuzzy-FishNet**. Fuzzy-FishNet’s strength lies in precision-recall but not so much in feature-selection stability. However, it should be noted that the RBNAs’ feature-stability may be inflated due to hypersensitivity.

Given that Fuzzy-FishNet complexes are enriched for highly significant SP-proteins, supported by the RBNAs, relatively stable and are rankable due to non-flat p distribution, this makes it a useful technique for feature selection in proteomics data.

### Significant features in proteomics and genomics data are poorly corroborative

To demonstrate that Fuzzy-FishNet also works on genomics data, analysis was repeated on the full TCGA dataset. Table 3 shows that the fisher p is low for the top 6 features while precision-recall performance is good relative to the other methods such as standard t-test and hypergeometric enrichment. Figure 5A shows that selected complexes are informative, and their constituent protein expression can clearly distinguish sample classes with little error (one misclassification). Comparing genomics complexes against proteomics complexes revealed that there is little overlap. Only 2 complexes overlapped, and there appeared to be few shared proteins amongst the significant complexes.

**Table 3.** **Top 5 complexes selected by Fuzzy-FishNet for renal cancer genomics data derived from TCGA.** The table shows that CORUM ID for the complex, the contingency table and the corresponding Fisher exact probability that was calculated.

| A | complex | class | hits | non-hits | fisher probability |
| --- | --- | --- | --- | --- | --- |
|  | 929 | cancer | 76 | 1119 | 2.18E-21 |
|  |  | normal | 246 | 1059 |  |
|  | 626 | cancer | 125 | 341 | 5.81E-21 |
|  |  | normal | 15 | 361 |  |
|  | 5589 | cancer | 0 | 217 | 1.21E-19 |
|  |  | normal | 64 | 191 |  |
|  | 4869 | cancer | 62 | 97 | 7.66E-19 |
|  |  | normal | 0 | 124 |  |
|  | 127 | cancer | 1 | 123 | 2.35E-14 |
|  |  | normal | 47 | 90 |  |
|  | 157 | cancer | 0 | 155 | 7.41E-12 |
|  |  | normal | 36 | 131 |  |

| B |  | precision | recall | F-score | sd_precision | sd_recall |
| --- | --- | --- | --- | --- | --- | --- |
|  | f_fishnet_4 | 0.759 | 0.761 | 0.760 | 0.081 | 0.081 |
|  | f_fishnet_6 | 0.797 | 0.798 | 0.798 | 0.074 | 0.074 |
|  | f_fishnet_8 | 0.826 | 0.826 | 0.826 | 0.071 | 0.071 |

| C |  | precision | recall | F-score | sd_precision | sd_recall |
| --- | --- | --- | --- | --- | --- | --- |
|  | sp_4 | 0.945 | 0.390 | 0.552 | 0.019 | 0.069 |
|  | sp_6 | 0.955 | 0.538 | 0.689 | 0.017 | 0.054 |
|  | sp_8 | 0.955 | 0.538 | 0.689 | 0.017 | 0.054 |
|  | hypgeo_4 | 0.938 | 0.303 | 0.458 | 0.038 | 0.100 |
|  | hypgeo_6 | 0.953 | 0.465 | 0.625 | 0.029 | 0.086 |
|  | hypgeo_8 | 0.953 | 0.465 | 0.625 | 0.029 | 0.086 |

**Figure 5.**
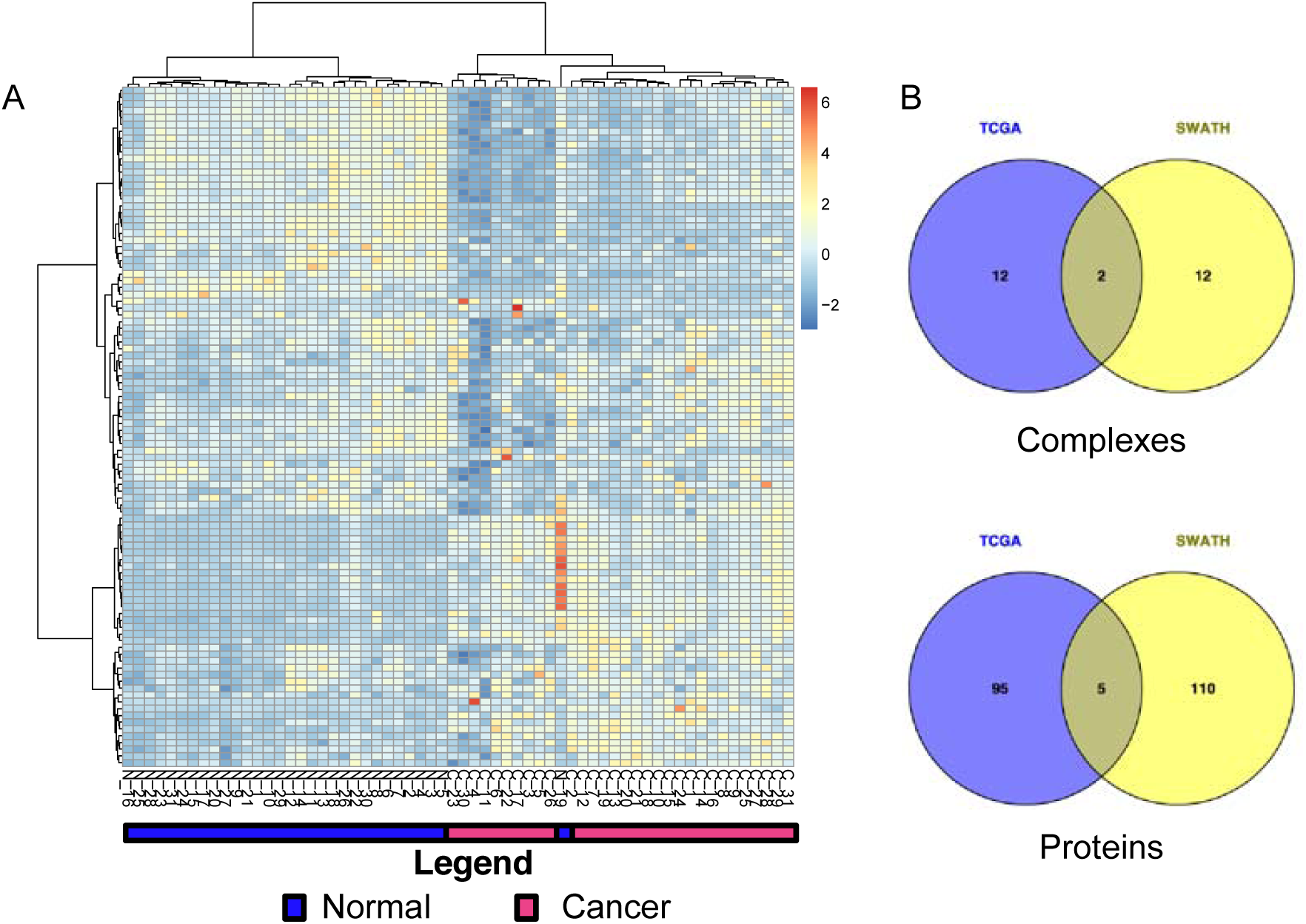
Fuzzy-FishNet also works on genomics data but selected features have little correspondence to proteomics. A: Fuzzy-FishNet can reliably separate sample classes based on the selected features. The tree (Euclidean Distance, Ward’s linkage) shows that using Fuzzy-FishNet, the sample classes can be reliably separated (with the exception of one normal sample). **B: Selected genomic features have poor correpondence to proteomics features.** The venn diagrams shows the complex overlap, and the corresponding protein overlap. Only 2 complexes, corresponding to 5 proteins, matched.

It is possible that the alpha genes for genomics data might be drastically different since many more genes are being considered relative to proteins in proteomic data. To counteract this possible effect, the genomics data matrix was subsetted to only include genes corresponding to proteins identified in the SWATH proteomics screen and the same analysis repeated.

With the subsetted TCGA matrix, Figure 6A shows improvements in the hierarchical clustering (no misclassifications were made this time). However, overlaps remain abysmal. Clearly this means that the top complexes selected by genomics and proteomics screens do not corroborate. It should also be noted that the 2 overlapping complexes in the subsetted genomics dataset are not the same. In fact, there are no overlaps amongst the top 14 complexes selected between the full and subsetted TCGA datasets. Furthermore, none of the overlapping TCGA (all and subset) selected complexes are amongst the top 5 in proteomics screen.

**Figure 6.**
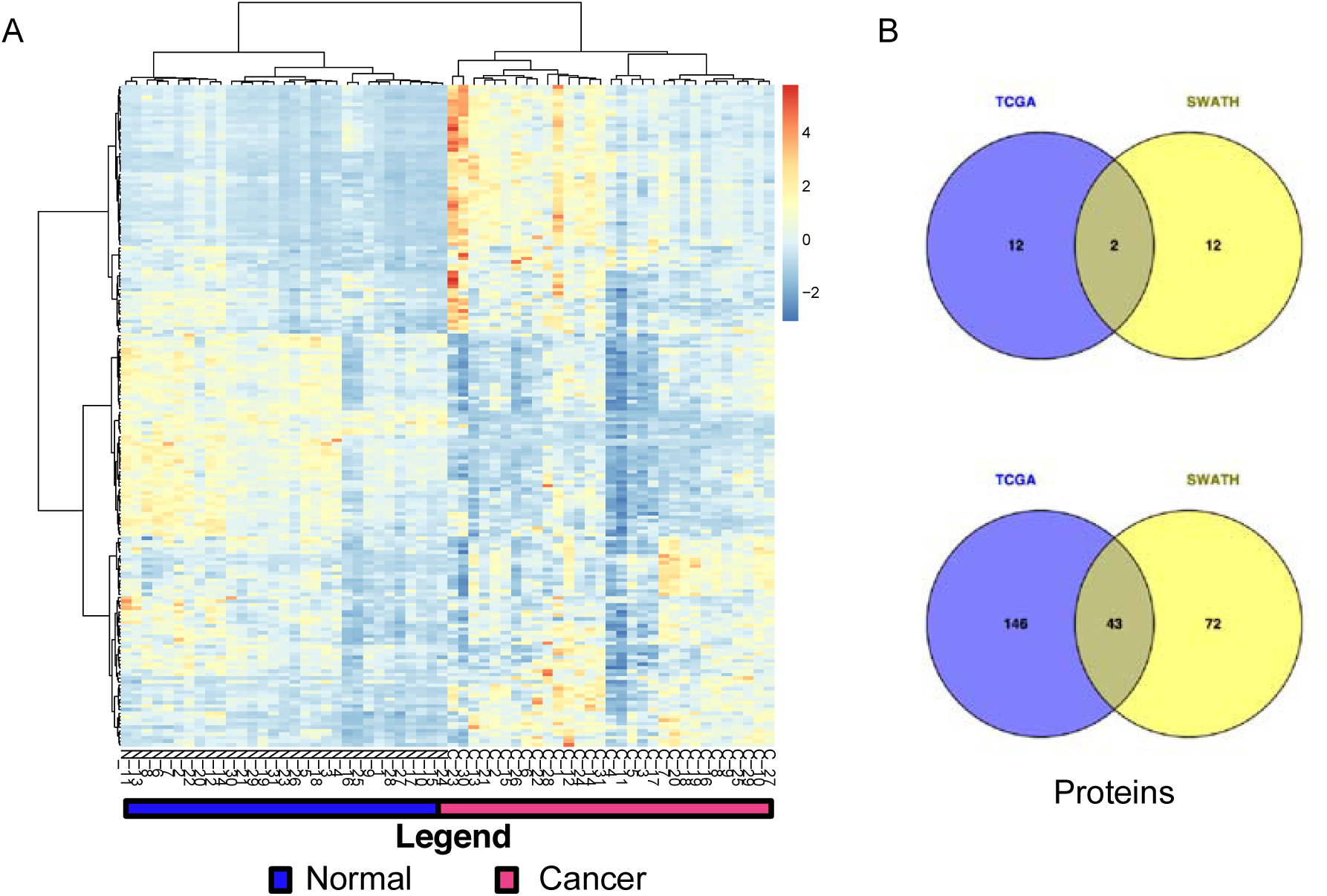
Subsetting genomics data to corresponding SWATH proteins did not improve proteomic-genomic correlation. A: Subsetted genomics data can still give rise to informative features. The heatmap (Euclidean Distance, Ward’s linkage) shows that using Fuzzy-FishNet on the subsetted genomics dataset (only include genes that corresponded to the proteins identified in SWATH), the sample classes can be reliably separated (interestingly, the clustering quality appeared better than using all genes; c.f. Figure 5A). **B: Post-subsetting, selected genomics features still have poor correlation to proteomics features**. The venn diagrams shows the complex overlap, and the corresponding protein overlap. 2 complexes, corresponding to 43 proteins, matched.

Proteomics and genomics measurements are known to be poorly-correlative (22-24). While it may be an attractive idea that networks can help to improve correlations between proteomics and genomics data (10), it appears that in practice the divide is not so easily bridged.

#### Integrating Fuzzy-FishNet with proteomics reveals a key role for mitochondrial complexes

The top three complexes consistent between FishNet and Fuzzy-Fishnet were the 55S ribosome (CORUM Complex ID 320), 28S ribosomal subunit (CORUM Complex ID 315) and F1/F0-ATP Synthetase (CORUM Complex ID 563). All three complexes are mitochondrial in origin. Based on the contingency tables in Figure 1A, all are overexpressed in the cancer class.

55S ribosome is involved in protein biosynthesis within the mitochondrial matrix (25). The 55S ribosome is composed of two subunits, the 28S subunit (which was also detected as overexpressed) and a 39S subunit. Not much has been reported on the mechanistic association of mitochondrial ribosomes with renal cancer. Increase in mitochondrial biogenesis is correlated to increased basal oxygen consumption, which in turn, leads to increased energy production. This phenomenon is well reported in Acute Myelogenous Leukemia (AML) (26). Skrtic *et al* showed that inhibiting mitochondria ribosome proteins using tigecycline as selectively killed leukemia stem and progenitor cells (26). Perhaps a similar strategy could also be deployed for renal cancer.

F1/F0-ATP Synthetase is involved in energy production using the oxidative phosphorylation pathways along the inner membrane of the mitochondrial wall (27). Recent work by Wang *et al* (28) also implicates the involvement of the oxidative phosphorylation pathways, and correlated this to metastatic outcome. To check if the high ranking of F1/F0-ATP Synthetase is strongly correlated with severe outcome, patients 2 and 8 (whom suffered from severe cancer) were removed from the data matrix and Fuzzy-FishNet repeated. F1/F0-ATP Synthetase dropped from rank 3 to 4 (p = 3.85e-04), which suggests the association of F1/F0-ATP Synthetase with severe renal cancer, at least based on our dataset, is limited. On the contrary, the 28S subunit suffered a pronounced drop from rank 2 to 5 (p = 4.84e-04), which implies stronger association with the severe phenotype.

## Conclusions

Fuzzy-FishNET is a new addition to the RBNA arsenal, and excels in precision-recall while maintaining small feature selection set. Using clear cell renal cancer as a case study, we demonstrated that the technique works well on both genomics and proteomics data. Cross-platform comparative analysis using Fuzzy-FishNet however, shows that the gulf between proteomics and genomics is not easily bridged, and significant features stably selected in genomics seldom match its proteomics counterpart.

## Acknowledgements

WWBG is funded by a Professorship of Bioinformatics, School of Pharmaceutical Science and Technology, Tianjin University, China.

### Author contributions

WWBG designed, implemented the bioinformatics method and pipeline, performed analysis, and wrote the manuscript.

## Supplementary Figures

**Supplementary Figure 1.**
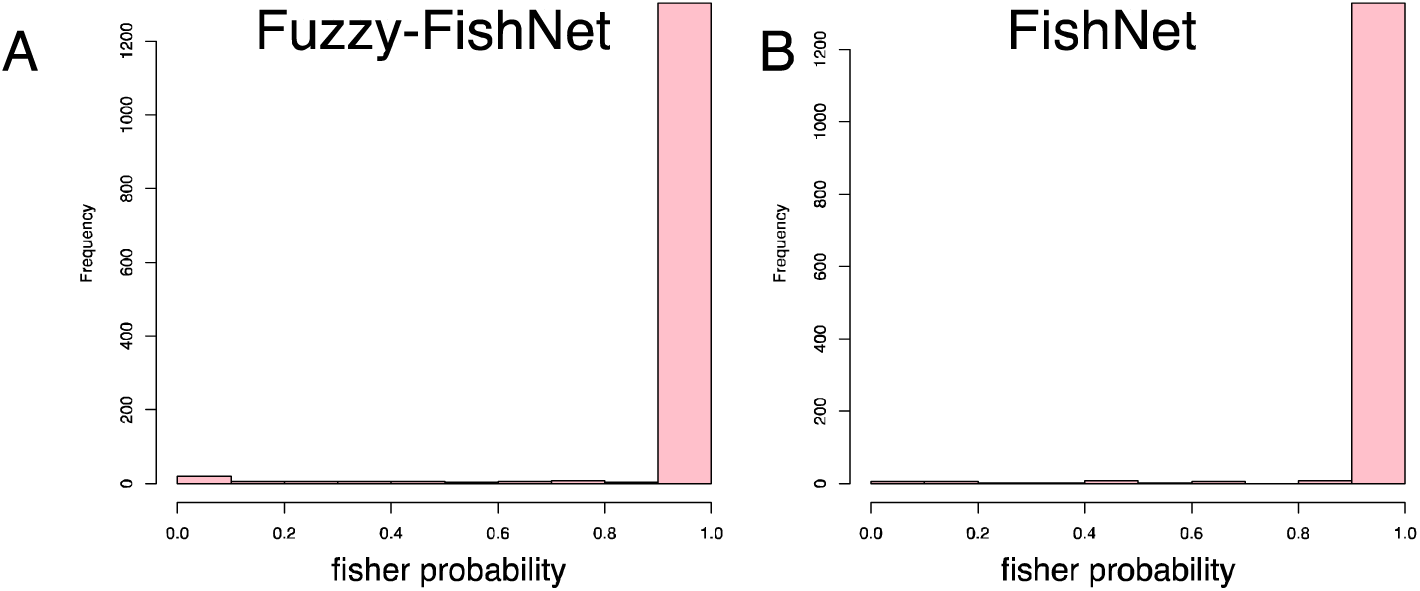
Distribution of fisher probabilities across 1363 protein complexes for Fuzzy-FishNet (A) and FishNet (B). Only a minority of complexes have small fisher exact probability. This shows that the calculation approach where we summed the signal across samples within classes does not lead to large selection sizes.

**Supplementary Figure 2.**
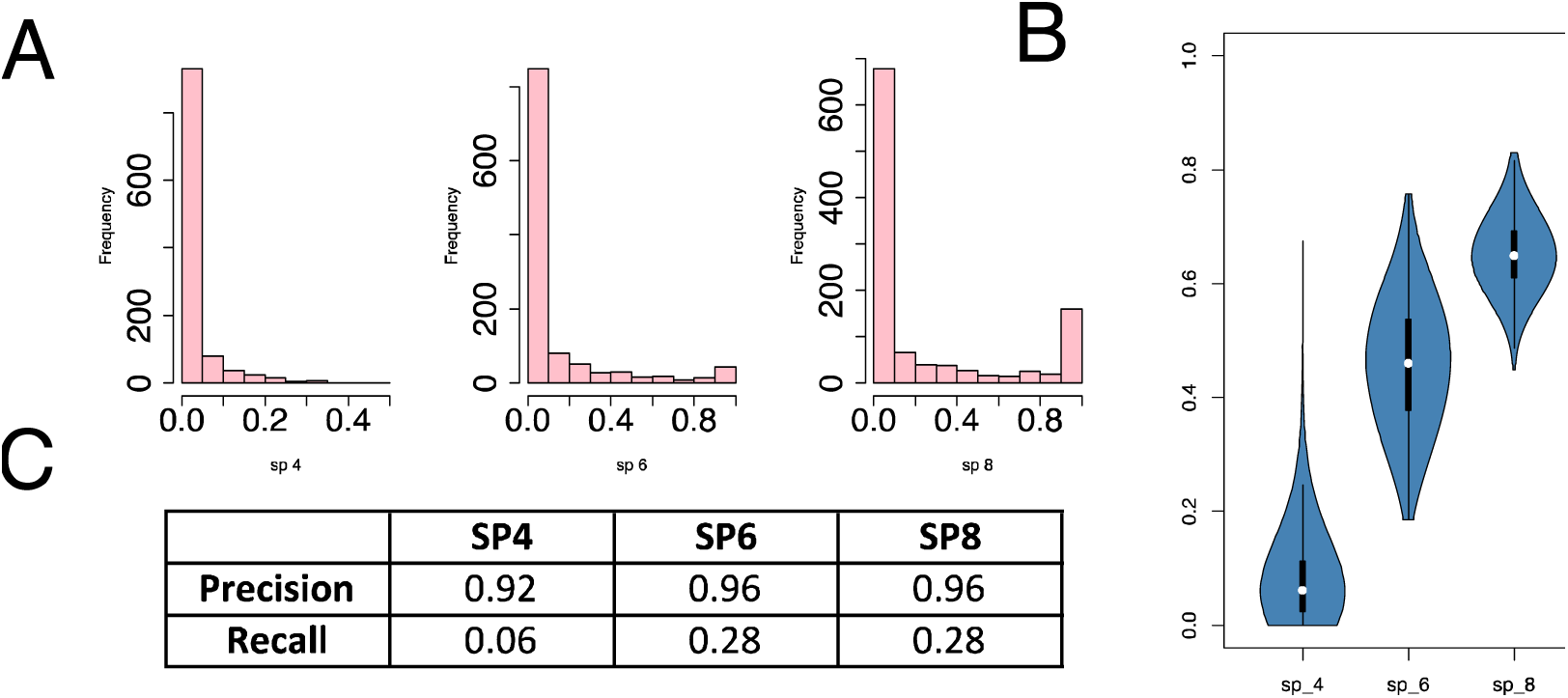
Evaluating single protein (SP) two sample t-test statistics A: Feature-stability following top 500 SP feature filtering. The proportion of stable features drops significantly when only the top 500 features per simulations are kept. **B: Pairwise feature-selection similarity following top 500 SP feature filtering**. Following feature filtering, the pairwise similarity decreases dramatically (y-axis, Jaccard coefficient) although it is still better than hypergeometric enrichment (HE). **C: Precision/recall following top 500 SP feature filtering**. After adjusting the critical value threshold to keep the top 500 features. Use of similar threshold on random subsamplings shows that precision is well maintained but recall drops further.

